# Gastric cancer treatment target identified from an accelerated Helicobacter-induced gastric cancer mouse model

**DOI:** 10.1101/2025.06.14.659641

**Authors:** Emma Mathilde Kurstjens, Kristin Cox, Prerna Bali, Siamak Amirfakhri, Jonathan Hernandez, Ivonne Lozano-Pope, Christopher Benner, Michael Bouvet, Marygorret Obonyo

**Author notes:** Corresponding author: Marygorret Obonyo, Department of Medicine, School of Medicine, University of California, San Diego 9500 Gilman Drive, MC 0640, La Jolla, CA 92093-0640, USA.

## Abstract

*Helicobacter pylori* (*H. pylori*) infection and consequent inflammation leads to gastric cancer (GC). Despite the prevalence of this bacterium and availability of genomic data, targeted therapies for GC are still early in development. Previously in our accelerated *Helicobacter*-induced gastric cancer mouse model we identified several differentially expressed genes (DEGs), including PSMB8 (proteasome subunit beta type 8, also called LMP7); one of the immune subunits of the immunoproteasome, which has been associated with disease severity in multiple cancers. We observed elevated expression of PSMB8 in our accelerated gastric cancer model, in the human gastric cancer cell line (MKN45), and in gastric cancer patient samples. Moreover, we identified carfilzomib as a potential drug that targets PSMB8. Therefore, to test its efficacy against gastric cancer, nude mice were subcutaneously implanted with MKN45 derived tumors and treated with carfilzomib, alone or in combination with 5-fluorouracil (5-FU), the standard care drug. The effectiveness of drug treatment was measured by tumor growth, cell proliferation, and apoptosis. We observed that carfilzomib retarded tumor growth, inhibited cell proliferation and induced apoptosis. These results strongly suggest that carfilzomib has a robust anti-tumor activity and is a suitable drug candidate for targeted therapy in gastric cancer.

## INTRODUCTION

*Helicobacter pylori* (*H. pylori*) infection and consequent inflammation leads to gastric cancer (GC). Based on recent meta-analysis, *H. pylori* infected 43.9% of adults and 35.1% of children and adolescents globally (1). *Helicobacter* has been classified as class I carcinogen and infection with this bacterium has been shown to increase the risk of developing GC by 1-3% (2, 3). GC is the third leading cause of cancer related deaths with an estimate of 770,000 deaths reported in 2020 (4, 5). Moreover, *H. pylori* caused over a third of infection-induced cancers in 2020, surpassing HPV, according to the World Health Organization (6). Despite the overwhelming prevalence of this bacterium and availability of genomic data, GC has a high mortality rate due to late diagnosis. The trend of late diagnosis is due to the non-specificity of common symptoms – dysphagia and weight loss- and their tendency to appear during the advance stages of disease (7, 8). Advanced stage diagnosis has a notable effect on 5-year survival rate, which is 36.4% in the US (9). Consequently, there is an unmet medical need for novel effective targeted therapies (8).

Targeted therapies for gastric cancer are still early in development. Since most cases of GC are diagnosed at stages III - IV, standard procedure of care consists of surgical resection and a nonspecific chemotherapy regimen called FLOT (5-fluorouracil, leucovorin, oxaliplatin, and docetaxel) (7,8). Thus, to address this gap in patient care, we previously identified a target protein in the TLR4 signaling pathway called MyD88 and hypothesized that removing this protein would reduce inflammation and hinder neoplasia; instead, we found that *Myd88^-/-^* mice infected with *Helicobacter felis* (*H. felis*; analogous to *H. pylori*) aggressively progressed towards gastric cancer, thus, now referred as the accelerated gastric cancer model (10, 11). Moreover, our RNA-seq analysis revealed several differentially expressed genes (DEGs) associated with severe disease pathology, such as indoleamine 2,3-dioxygenase 1 (*Ido1*), guanylate binding protein 2 (*Gbp2*), interferon regulatory factor 1 (Irf1), beta-2 microglobulin (*B2M*), and proteasome subunit beta 8 (*PSMB8*) (11, 12).

Immunoproteasomes consist of two outer α- and two β-subunits. They have a vital function within the cell where they maintain cellular protein homeostasis and process antigens for presentation via the major histocompatibility complex upon stimulation. Upon stimulation the *β1*, *β2* and *β5* subunits are substituted by corresponding immune subunits *β1i* (PSMB9), *β2i* (PSMB10) and *β5i* (PSMB8) (13, 14, 15). Cancer cells rely heavily upon their function for survival and proliferation (13, 15, 16). Moreover, upregulation of immunoproteasomes has been observed in blood cancers such as multiple myeloma, as well as multiple different solid tumors (13). Thus, suggesting that immunoproteasomes can serve as targets in treating cancer.

Therefore, in this study we first evaluated the expression of the immune subunits of the immunoproteasome, PSMB8, 9, and 10 in our accelerated model. Then we investigated their expression in the human gastric cancer cell line (MKN45) and observed PSMB8 being the highest expressed gene. Thereafter, we validated our results in the human gastric cancer patient tissue samples to assess its relevancy in GC. We found carfilzomib as a potential drug that targets PSMB8 specifically and studied its efficacy in treatment of GC either alone or in combination with 5-fluorouracil (5-FU)-a standard treatment drug-by monitoring tumor growth, cell proliferation and apoptosis. The data from the present study indicated that carfilzomib hampered tumor growth and cell proliferation and induced apoptosis, making it a promising drug for targeted treatment in gastric cancer.

## MATERIALS AND METHODS

### RNA extraction

RNA was extracted from subcutaneously implanted MKN45 tumors or human gastric tissue biopsy samples obtained from the Biorepository, University of California, San Diego. Gastric tissue samples were frozen and pulverized using a chilled mortar and pestle, then homogenized in 1 ml of TRIzol reagent (Invitrogen, Carlsbad, CA) with a Dounce homogenizer. Thereafter, RNA was extracted using Direct-zol RNA MiniPrep Kit (Zymo Research, Irvine, CA) as per the manufacturer’s instructions and stored at -80°C until further use.

### cDNA synthesis and quantitative real-time RT-PCR

2µg of RNA isolated from either human tumor samples or subcutaneously implanted MKN45 tumors was reverse transcribed into cDNA using the High Capacity cDNA Reverse Transcription Kit (Thermo Fisher, Waltham, MA) as per manufacturer instructions. Real-time RT-qPCR (reverse transcription quantitative polymerase chain reaction) was performed on the StepOne Plus Real Time PCR system (Applied Biosystems, Carlsbad, CA) using SYBR Green Supermix (Biorad, Irvine, CA). 1μl of cDNA was used per well for a total of 10μl reaction mix. The amplification conditions were as follows: initial cycle of 95°C for 5 min, annealing at 60°C for 20 sec, and extension at 72°C for 40 sec. Expression levels of PSMB8, PSMB9, and PSMB10 were normalized to the housekeeping gene HPRT-1. The data collected were analyzed using comparative cycle threshold calculations and plotted using GraphPad Prism software (La Jolla, CA, USA). The primers used are listed in Table S1.

### Drug Identification

Carfilzomib was identified using a public pharmaceutical database called the Drug Gene Interaction database (DGIbd) (18).

### Animals

All the animal procedures were approved by the University of California San Diego Institutional Animal Care and Use Committee (IACUC) and conducted following accepted veterinary standards and ARRIVE guidelines. Athymic male nude mice, aged 4-6 weeks, were used for the experiments. The animals were fed an autoclaved diet and housed in a barrier facility. Prior to any surgical procedure, the mice were anesthetized with a solution of xylazine, ketamine, and phosphate-buffered saline (PBS) via intraperitoneal injection. At the conclusion of the study, mice were anesthetized with isoflurane and euthanized by cervical dislocation.

### Xenograft establishment

A poorly differentiated human gastric cancer cell line, MKN45, was used for these experiments. Cells were cultured in Roswell Park Memorial Institute media (RPMI) with 10% fetal bovine serum (FBS) and 1% penicillin. 1 x 10^6^ MKN45 cells suspended in 100 µL of PBS were injected into the bilateral flanks and shoulders of nude mice to initially establish subcutaneous models. Once subcutaneous tumors grew to approximately 1 cm, subsequent passages were performed by harvesting 1 mm^3^ fragments and implanting them into additional nude mice. For the experiments, a ∼5 mm incision was made on the mid-back of nude mice and a single 1 mm^3^ tumor fragment was implanted into the right flank. The incision was closed with a simple interrupted 6-0 vicryl suture (Ethicon Inc.). Tumors were allowed to grow for 3 weeks before mice received any treatment. This protocol is summarized in Figure 1.

**Figure 1:**
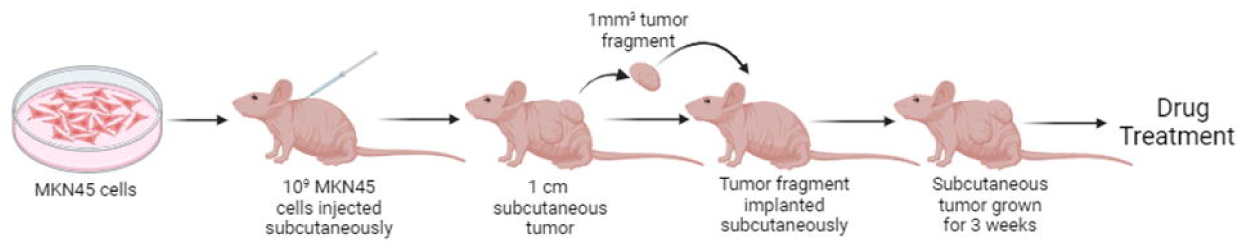
Schematic diagram showing the process of establishing the xenograft model in preparation for experimental treatment.

### Drug Dosing

5-Fluorouracil (5-FU) was purchased from Acros Organics BVBA and resuspended in PBS to make a stock solution of 10 mg / mL. Carfilzomib was purchased from Onyx Pharmaceuticals and resuspended in 2 mL of DMSO to make a stock solution of 50 mg / mL. 5-FU and carfilzomib were administered at a dosage of 50 mg / kg, and 5 mg / kg, respectively. PBS containing 2% DMSO served as the placebo control.

### Treatment Regime

Nude mice were divided into 4 groups (n=10/group) - 5-FU (administered 5- FU only), Carfilzomib (administered carfilzomib only), Combination (administered both 5-FU and carfilzomib) and Control (administered placebo control). Both drugs and the placebo control were administered via intraperitoneal (IP) injections with a total volume of 150 µL. One dose of 5-FU was administered per week for 8 weeks, and Carfilzomib and the placebo control were administered for consecutive two days per week for 8 weeks. The schedule is summarized in Figure 2.

**Figure 2:**
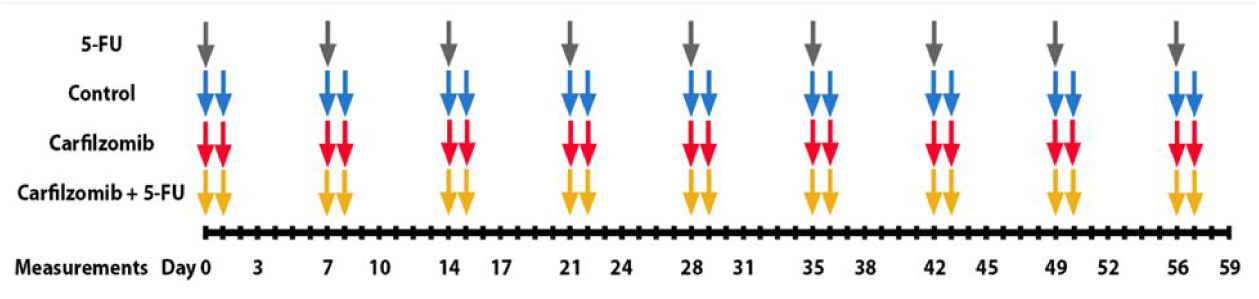
Schematic diagram showing the schedule of intraperitoneal injections delivered to experimental mice.

### Monitoring of Mice

Mice were monitored for both tumor growth and body weight twice a week. Tumor size was measured using calipers in both the width (W) and length (L) dimensions. Tumor volume was then calculated using the following formula, (W x W x L) / 2. The weights of the mice were taken using a scale that was zeroed prior to each use.

### Criteria for Termination

The following criteria were used to determine when mice should be euthanized in order to comply with IACUC standards; 1) Tumor size > 1.5 cm in any dimension, 2) Ulceration of tumor through skin surface, 3) Weight loss of 20%, 4) Signs of physical distress such as hunched posture, lethargy, or inability to ambulate. Termination due to the fourth criterion was not observed in this study. 11 mice were euthanized prior to the conclusion of the monitoring period. In the control group, 4 mice were euthanized (3 for size and 1 for ulceration). In the 5-FU group, 2 mice were euthanized for tumor size. In the Carfilzomib group, 3 mice were euthanized (1 for size and 2 for ulceration). In the combination group that received both 5-FU and Carfilzomib, 2 were euthanized Carfilzomib (1 for size and 1 for weight loss).

### Paraffin Embedding of Tumor Tissue Samples

Subcutaneous human cell line (MKN45) derived tumors were paraffin embedded according to the following procedures. A portion of tumor tissue from each mouse was fixed in neutral buffered 10% formalin, then embedded in paraffin and sectioned on a microtome. 5 μm sections were mounted onto glass slides and then either subjected to TUNEL assay or immunohistochemistry (IHC).

### TUNEL Assay

TUNEL (terminal nucleotidyl transferase-mediated dUTP-biotin nick end-labelling) assay was performed on paraffin embedded gastric tumor tissues slides using the ApopTag Peroxidase In Situ Apoptosis Detection Kit (Sigma Aldrich, St. Louis, MO) following manufacturer’s instructions with two modifications, wherein peroxidase substrate was applied for 30 seconds and methyl green was applied for 5 minutes. The quantification of positive staining was performed via imaging with the Olympus VS200 Slide Scanner (access provided by UCSD School of Medicine Microscopy Core, grant # NS047101) followed by analysis with QuPath software (version 8.2.0).

### Immunohistochemistry (IHC)

IHC was performed with rabbit anti-human Ki67 primary antibody (ab16667, Abcam, Cambridge, UK). Paraffin embedded gastric tissue slides were deparaffinized in xylene and decreasing ethanol dilution series, followed by antigen retrieval in citrate buffer heated to 90°C for 20 minutes. The sections were then washed in 0.3% Triton X-100 solution and blocked for 1 hour with a solution of 3% BSA, 0.1% Tween 20, 0.1% Triton X-100, and 5% normal goat serum in PBS at 22 – 25 °C. The sections were washed and stained with primary antibody, 1:100 dilution, at 4°C overnight. This is followed by incubation with an HRP-conjugated anti-rabbit secondary antibody (Cell Signaling, cat#7074S), 1:200 dilution at 22 – 25 °C for 1 hour. Peroxidase substrate was applied for 5 minutes, and Harris’ hematoxylin was applied for 2 minutes. The quantification of positive staining was performed via imaging with the Olympus VS200 Slide Scanner (access provided by UCSD School of Medicine Microscopy Core, grant # NS047101) followed by analysis with QuPath software (version 8.2.0).

### Quantification and statistical analysis

Statistical analysis was performed using GraphPad Prism (La Jolla, CA, USA). ANOVA with Bonferroni’s correction (for normal distribution) were used for multiple comparisons. P-values ˂0.05 were considered statistically significant.

## RESULTS

### Elevated PSMB8 expression observed in accelerated murine gastric cancer model

Previously, in our RNA-Seq data we observed high expression of certain interferon stimulated genes (ISGs) in our accelerated model of gastric cancer-designated as differentially expressed genes (DEGs) (11,12) which included PSMB8, an immune subunit of immunoproteosome, therefore we evaluated the expression of the immunoproteosome immune subunits (PSMB8, 9 and 10) in our accelerated model. We observed high expression of all the genes in the accelerated model at both 25-week and 47-week than in the standard model (infected WT) as seen in Figure 3. This increased expression in the accelerated model may suggest a potential role of these genes in disease progression.

**Figure 3:**
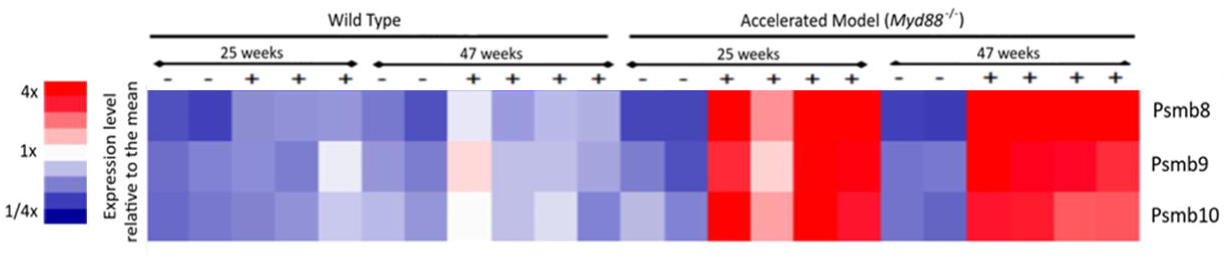
Heatmap showing expression of Immune subunits of Immunoproteosome in standard and accelerated models. Relative expression of PSMB8, PSMB9 and PSMB10 was measured in standard and accelerated models post infection with *H. felis* at 25 weeks and 47 weeks. Indicated by the color scale on the left, a brighter red color indicates a greater degree of relative expression as compared to the mean, while a deeper blue indicates a lesser degree of expression as compared to the mean. Positive ‘+’ and negative ‘-’ symbols indicate the infection status of each mouse assayed.

### Elevated PSMB8 expression observed in MKN45 cells and gastric cancer patient tissue samples

Since we observed high expression levels of PSMB8, PSMB9, and PSMB10 in our accelerated model therefore we wanted to examine their expression levels in MKN45 cells, a human gastric cell line via RT-qPCR. Highly elevated levels of PSMB8 were observed in comparison to PSMB9 and PSMB10 (Figure 4). Thus to further validate our findings in the MKN45 cell line we measured the expression levels of PSMB8 in human gastric cancer tissue samples. High expression of PSMB8 was observed (Figure 5). Furthermore, in the paired tissue sample, the expression of PSMB8 was observed to be 137.6 fold higher in the tumor sample than the normal sample (Figure 5). Thus, confirming the relevance of the study and use of PSMB8 as a potential drug target candidate.

**Figure 4:**
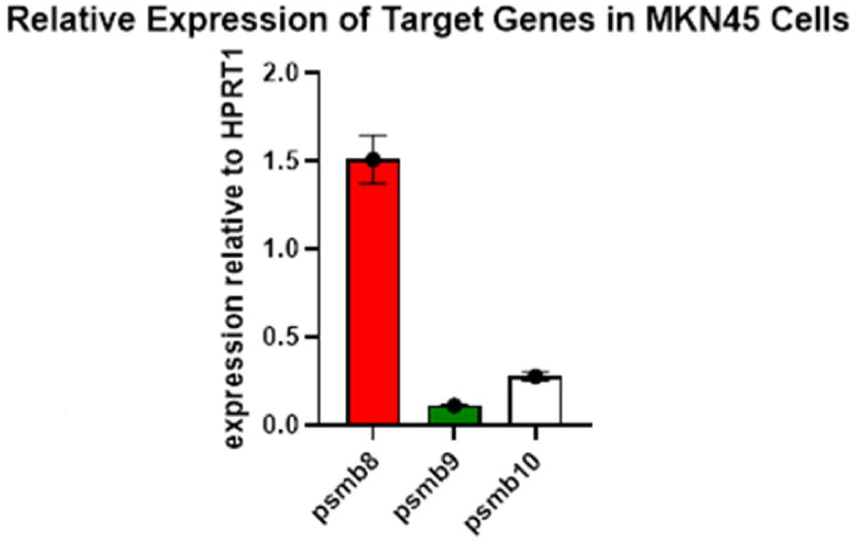
Relative expression of select DEGs in MKN-45 cells. RT-qPCR was performed with cDNA from MKN45 cells to detect levels of PSMB8, PSMB9, and PSMB10. Expression is measured via relative to expression of housekeeping gene HPRT1. Data are represented as mean ± SEM.

**Figure 5:**
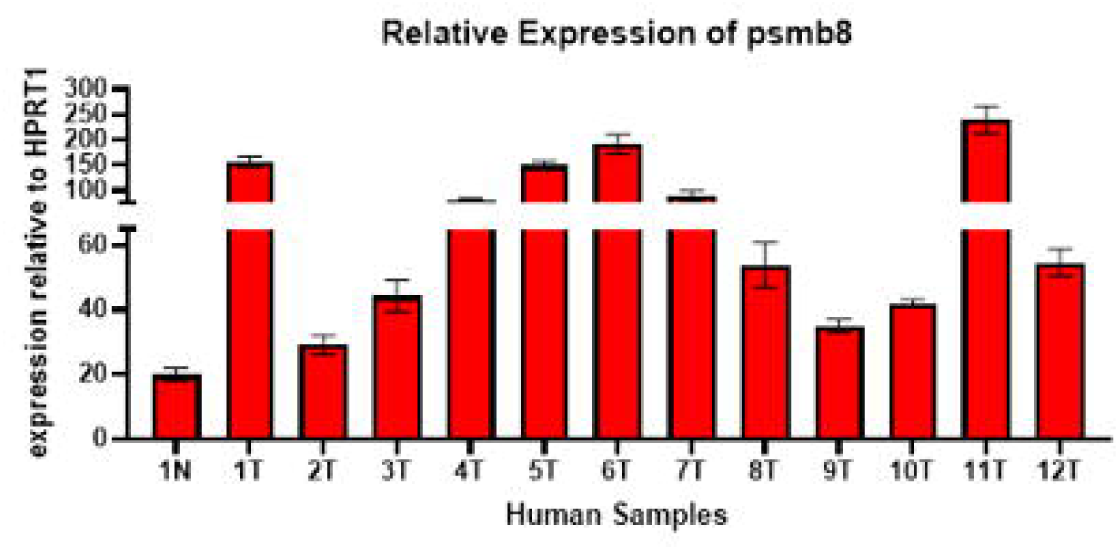
PSMB8 expression in gastric cancer patient samples. RT-qPCR was performed with cDNA from GC patients (n=12) to detect levels of PSMB8. Individual ‘1’ has a paired sample of normal tissue, signified by the ‘N’ at the end of the label. All sample labels ending with ‘T’ indicates a tumor sample. Expression is measured via relative to expression of housekeeping gene HPRT1. Data are represented as mean ± SEM.

### Carfilzomib treatment significantly slows tumor growth

Therefore, to further assess the potential of PSMB8 as a drug target we searched for a public pharmaceutical database for identifying a drug that targets PSMB8 and identified carfilzomib as a potential drug. We then evaluated its efficacy by monitoring tumor growth, cell proliferation and apoptosis, as a potential drug for treatment of gastric cancer in our xenograft murine model.

The increase in tumor volume was monitored over the course of the treatment period of 59 days. The mice in the control group showed a rapid increase in tumor volume over the given time period than the treatment groups (5FU, carfilzomib, and the combination group). The mice treated with carfilzomib showed significantly lower rates of tumor growth than the control at two time-points: (p= 0.0477392) at 38 days and (p= 0.0350377) at 59 days (Figure 6). Within the treatment groups the Carfilzomib treatment group showed slow increase in tumor volume as compared to the other two treatment groups (Figure 6). These results suggest that carfilzomib retards tumor growth, and strengthens it potential use in treatment of gastric cancer.

**Figure 6:**
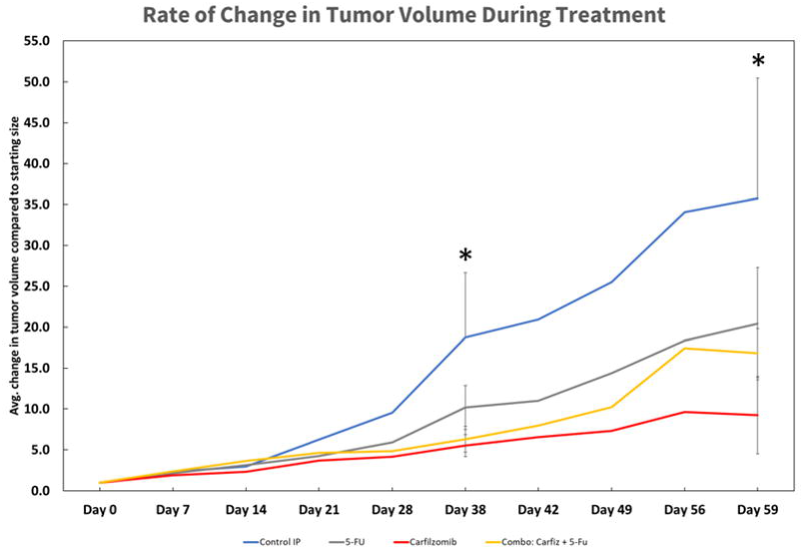
Tumor Volume Measurements. Rate of increase in tumor volume was measured over the span of 59 days. Asterisks (*) indicate the statistically significant difference between the data points on the blue line, indicating the progress of control mice, and the data points on the red line, indicating carfilzomib treated mice. Data are represented as mean ± SEM.

### Carfilzomib induces apoptosis

We performed TUNEL assay to measure apoptosis in the different treatment groups. TUNEL (terminal nucleotidyl transferase-mediated dUTP-biotin nick end-labelling) staining tags the double stranded DNA breaks that occur during apoptosis, thus indicating which cells have been affected by treatment. We observed that the combination of carfilzomib and 5FU shows a significant increase in positive TUNEL staining as compared to 5FU alone (p = 0.0266) and control (p = 0.00318) (Figure 7A). A marked increase in positively stained cells was observed in the carfilzomib and combination treatment groups as seen in the representative images (Figure 7B). Therefore, the increase in positive TUNEL staining suggests that carfilzomib is inducing apoptosis and strongly supporting its potential use in treatment of gastric cancer.

**Figure 7:**
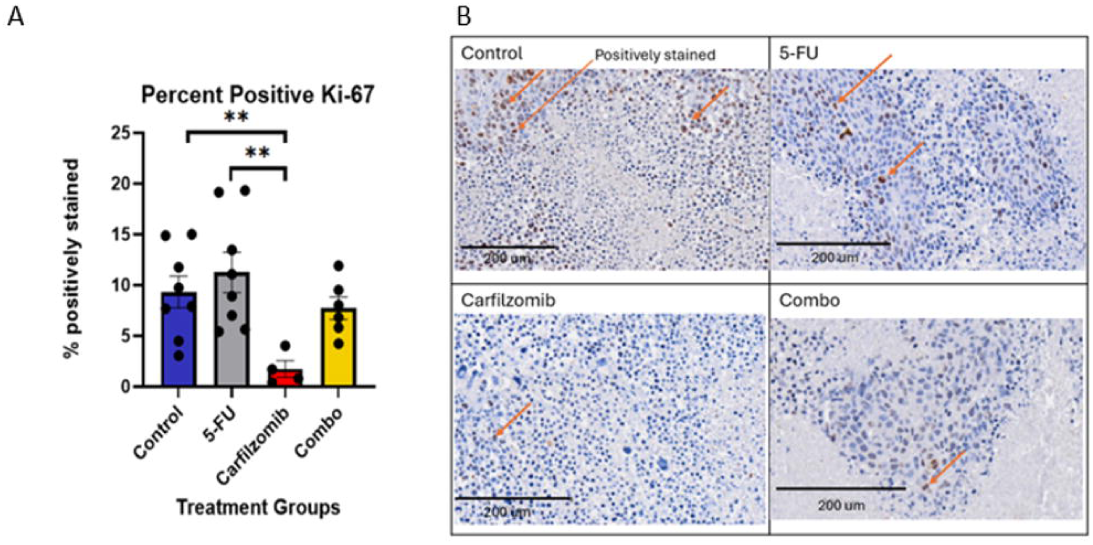
Quantification of Ki67 staining. A) A bar graph depicting the average percentage of positively stained cells for each group; the asterisks (*) indicate statistically significant differences. Data are represented as mean ± SEM.. B) Representative image of each group; orange arrows indicate a positive Ki67 stain. *, P value <0.05;**, P value <0.01; ***, P value <0.001.

### Cell proliferation significantly impeded by treatment with carfilzomib

Ki67 staining helps in quantifying the amount of cellular proliferation that has occurred during treatment. Inhibition of cancer cell proliferation is an indicative of an effective treatment. We observed that the carfilzomib treated group showed a significantly lower percentage of Ki67 positively stained cells compared to both the control (p=0.00847) and 5-FU (p=0.00838) groups (Figure 8A). A marked reduction in positive cells was observed in carfilzomib treatment group in comparison to the other groups (Figure 8B). This reduction in cell proliferation is indicative of successful treatment by carfilzomib.

**Figure 8:**
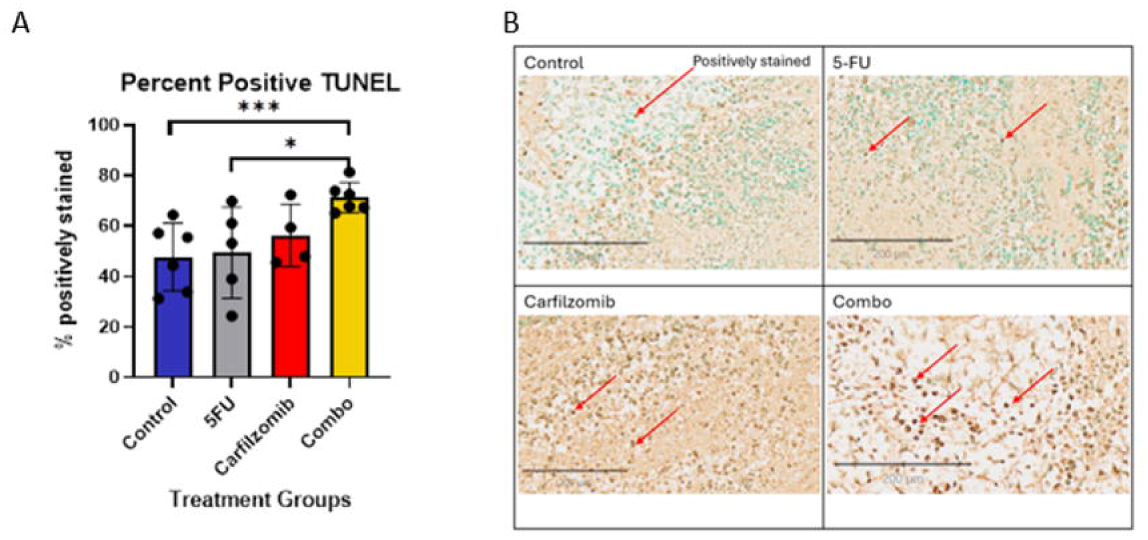
Quantification of TUNEL staining. A) A bar graph depicting the average precent of positively stained cells for each group; the asterisks (*) indicate statistically significant differences. Data are represented as mean ± SEM. B) Representative image of each group; orange arrows indicate a positive TUNEL stain. *, P value <0.05;**, P value <0.01; ***, P value <0.001.

## DISCUSSION

Our accelerated model revealed high expression of PSMB8 (10, 11) which we confirmed in both the human gastric cancer cell line MKN45 and patient samples via qPCR. Previous studies have shown that PSMB8 plays an important role in hepatocellular carcinoma through interactions with zinc finger family proteins (19). More broadly, a pan-cancer analysis found that of the 33 cancer types assessed, overexpression of PSMB8 was related to poor clinical outcomes (20). It has also been specifically noted that elevated nuclear expression of PSMB8 in gastric cancer patients has been correlated to a decrease in overall patient survival (21). Thus, making PSMB8 an ideal drug target candidate.

Repurposing drugs aids in developing a treatment regime faster than traditional drug development and is more cost effective (22, 23). Thus, after exploring the publicly available pharmaceutical databases, we identified carfilzomib as a potential drug capable of targeting PSMB8 (18). Carfilzomib, a second-generation proteasome inhibitor, was previously used to treat multiple myeloma, non-Hodgkin’s lymphoma, and leukemia (24, 25, 26). An epoxyketone that inhibits immunoproteasome activity by mimicking the structure of epoxomicin (25, 26). This leads to its binding to immunoproteasome catalytic subunit β5i encoded by PSMB8 with high specificity *in cellulo* selectively and irreversibly (24, 26), inducing apoptosis via both intrinsic and extrinsic pathways (27). Carfilzomib has been already approved by the FDA for treatment of patients with relapsing or refractory multiple myeloma in combination with dexamethasone (24, 28). Since it is an already approved drug, its usage as a drug for targeted therapy in gastric cancer can be permitted quickly (22, 23). Separately, it has been shown previously that selective and simultaneous inhibition of proteasome and immunoproteasome subunits, β5 and β5i, by carfilzomib induces a strong anti-tumor response in multiple myeloma in comparison to current available chemotherapies, such as bortezomib (26, 27). Hence making carfilzomib a suitable candidate for drug repurposing to treat gastric cancer.

Cancer is often treated with multiple drugs, with the goal of producing a greater, additive effect that a single drug could not achieve while minimizing toxicity to noncancerous cells, which can be aided by the application of a targeted treatment such as carfilzomib (14, 28, 29). The drug 5-FU is a primary component of FLOT, the standard treatment regime for gastric cancer patients. FLOT is a nonspecific chemotherapy regimen which has a known toxic effect on its patients (30). Therefore, to test carfilzomib’ s full treatment potential, subcutaneous tumor models were produced using an established gastric cancer cell line and treated with drugs 5-FU, carfilzomib, or a combination of both, in an effort to see if an additive effect could be achieved. We observed significant difference in average tumor volume between the placebo control group and the carfilzomib treated group. A previous study showed that oral recombinant methionase (o-rMETase) in combination with 5FU effectively suppressed tumor growth compared to the monotherapy treatment groups and untreated control (31). In contrary to these findings we observed that amongst the treatment groups the carfilzomib treatment group inhibited tumor growth to the greatest extent, indicating a relatively successful treatment.

Moreover, it is important to note that we observed a significant difference in tumor growth between placebo control and carfilzomib treatment group at day 38 and day 59, the final time point, attributing to the fact that treatment with carfilzomib limited the growth of the tumor. Thus, suggesting that treatment with carfilzomib may improve the clinical outcomes for patients with gastric cancer as it may help in retarding the tumor growth, thereby allowing an effective surgical resection. Previously it has been shown that treatment with FLOT preoperatively increased patient survival from a median of 35 months to 50 months as compared to older therapies; though statistically being non-significant (30). In our study carfilzomib monotherapy retarded tumor growth effectively followed by the combination treatment group in comparison to 5-FU monotherapy. Therefore, suggesting that treatment with carfilzomib preoperatively may increase the overall patient survival. Thereby strengthening its potential as a drug for treatment of patients with gastric cancer.

Apoptosis is a common measure of successful drug treatment, as it indicates effective killing of cancer cells. In this study we observed high induction of apoptosis in the combination and carfilzomib treatment groups, but the highest induction was observed in the combination treatment group which is similar to the findings observed by Li et.al, where they showed that treatment with a combination of TRAIL and 5-FU also induced significant apoptosis (32). Thus, suggesting that carfilzomib not only possesses an anti-tumor activity but also enhances the anti-tumor activity of 5-FU in combination.

Inhibition of cell proliferation is an indicator of anti-tumor activity measured by Ki-67 expression. Several studies have associated Ki-67 expression with effectiveness of chemotherapy in gastric cancer (33, 34), as previously shown in a study where high expression of Ki-67 was associated with shorter disease-free survival time and overall survival in gastric cancer patients who received neoadjuvant FLOT chemotherapy (34). Miyake et al. showed that combination of 5FU and o-METase showed reduced expression of Ki-67 (31). However, in our study we observed highest reduction in Ki-67 expression in carfilzomib treatment group compared to all the other treatment groups and control group. Therefore, suggesting that carfilzomib alone can effectively inhibit cell proliferation in tumor cells thus further supporting its use in treatment of gastric cancer.

This study has its own set of limitations. One limitation of this study is that the degree of inhibition of PSMB8 by carfilzomib was not measured *in vivo*. Another limitation being that the toxicity levels of this drug was not tested.

Further studies need to be done to understand the underlying molecular mechanism in carfilzomib treatment.

In conclusion, we suggest that carfilzomib is a promising drug candidate for targeted therapy. It possesses a robust anti-tumor activity as it is able to inhibit tumor growth by inducing tumor cell loss via apoptosis and impeding cell proliferation. Thus, showing a great potential of being a part of a targeted treatment plan for gastric cancer either in combination with the standard treatment plan already in place or a stand-alone monotherapy which may improve the overall disease free survival rate in patients with gastric cancer. However, further clinical evaluation in gastric cancer is a requisite to completely assess its true potential.

## ACKNOWLEDGMENTS

The present study was supported by the Department of Defense (DOD), under award W81XWH-20-1-0675.

## AUTHOR CONTRIBUTIONS

MO supervised and designed the study concept and research; CB supervised data analysis; EMK, JH, and ILP performed molecular work under the supervision of PB and MO; KC and SA performed xenograft mouse model and collected mouse samples under the supervision of MB; PB collected human gastric cancer biopsy samples; EMK performed data analysis; Manuscript is written by EMK, and PB. MO revised the manuscript. All authors read and approved the manuscript.

