## Supplementary material for "Gastric cancer treatment target identified from an accelerated Helicobacter-induced gastric cancer mouse model": Table S1

### SUPPLEMENTARY MATERIALS

**Table S1**: List of primers used in RT-qPCR.

| Gene Name | Forward/ Reverse |
|---|---|
| human PSMB8 | CCTTACCTGCTTGGCACCATGT/<br>TTGGAGGCTGCCGACACTGAAA |
| human PSMB9 | CGAGAGGACTTGTCTGCACATC/<br>CACCAATGGCAAAAGGCTGTCG |
| human PSMB10 | GGACAAGAGCTGCGAGAAGATC/<br>ATCTTGGACGCCACCATCCGTG |
| human HPRT1 | CCTGGCGTCGTGATTAGTGAT/<br>AGACGTTCAGTCCTGTCCATAA |
